# Do Synthesis Centers Synthesize? A semantic analysis of diversity and performance

**DOI:** 10.1101/518605

**Authors:** Edward J. Hackett, Erin Leahey, John N. Parker, Ismael Rafols, Stephanie Hampton, Ugo Corte, John M. Drake, Bart Penders, Laura Sheble, Niki Vermeulen, Todd Vision

## Abstract

Synthesis centers are a recently-developed form of scientific organization that catalyzes and supports a form of interdisciplinary research that integrates diverse theories, methods and data across spatial or temporal scales, scientific phenomena, and forms of expertise to increase the generality, parsimony, applicability, or empirical soundness of scientific explanations. Research has shown the synthesis working group to be a distinctive form of scientific collaboration that reliably produces consequential, high-impact publications, but no one has asked: do synthesis working groups produce publications that are substantially more diverse than those produced outside of synthesis centers, and if so, how and with what effects? We have investigated these questions through a novel textual analysis. We found that if diversity is measured solely by mean difference in the Rao-Stirling (aggregate) measure of diversity, then the answer is no. But synthesis center papers have significantly greater variety and balance, but significantly lower disparity, than papers in the reference corpus. Synthesis center influence is mediated by the greater size of synthesis center collaborations (numbers of authors, distinct institutions, and references) but even when taking size into account, there is a persistent direct effect: synthesis center papers have significantly greater variety and balance, but less disparity, than papers in the reference corpus. We conclude by inviting further exploration of what this novel textual analysis approach might reveal about interdisciplinary research and by offering some practical implications of our results.

---

Interdisciplinary research, widely heralded as a way to solve complex societal problems and to produce deeply original, even transformative, scientific knowledge, has been pursued and promoted for decades by scientists and science policymakers (NRC 2004; Porter et al. 2006; NSB 2008; Frodeman et al., 2010).^1^ Hopes that interdisciplinary research would arise through natural processes of blind variation and selective retention (Campbell, 1960), consilience (Wilson 1998), or convergence (Sharp 2011) are accompanied by interventions that create organizations and processes to foster interdisciplinary collaboration (see Palmer et al. 2016).^2^ Synthesis centers are among the most visible and potentially effective of such organizations. Beginning with the National Center for Ecological Analysis and Synthesis in 1995, the US National Science Foundation has invested in a series of synthesis centers culminating in the Socio-Ecological Synthesis Center (SESync).^3^ With their prominence, scale, and apparent success, synthesis centers have been strategic sites for several studies of the process and outcomes of their distinctive form of interdisciplinary collaboration (Hackett et al. 2008; Rhoten and Parker 2003; Hampton and Parker 2011; Hackett and Parker 2016), but no study yet has asked, *Do synthesis centers synthesize?* That is, if such centers integrate diverse concepts, theories, tools, techniques and data, then the publications of synthesis-center collaborations should be more diverse and more visible than other publications. We address these questions through semantic analysis of the text (e.g., titles, abstracts, and keywords) of published journal articles to compare the topical diversity of publications originating in synthesis centers with publications in a reference corpus of scientific literature.^4^

## What Is Synthesis?

Scientific synthesis is a form of interdisciplinary research that integrates diverse theories, methods and data across spatial or temporal scales, scientific phenomena, and forms of expertise to increase the generality, parsimony, applicability, or empirical soundness of scientific explanations (Carpenter et al., 2009; Hackett and Parker 2016). Synthesis occurs through collaboration among disciplinary or transdisciplinary experts, and therefore encompasses and extends beyond more typical forms of interdisciplinary research. Synthesis counterbalances scientific specialization, capitalizes on existing data, and addresses complex problems (Hackett et al. 2008; Palmer et al. 2016). When successful, synthesis draws specialties or disciplines together in novel configurations that open new spheres of inquiry and address societal challenges in original and effective ways (Carpenter et al. 2009; Baron et al. 2017; Wyborn et al. 2018).

The first synthesis center, the National Center for Ecological Analysis and Synthesis (NCEAS), was founded in 1995, funded by the US National Science Foundation and the State of California. NCEAS was designed to promote interdisciplinary collaborations that extended across academic disciplines and, in some cases, also included environmental policymakers and government officials to address problems of scientific and societal importance (Hackett et al., 2008). In doing so the center would also transform the practice and outcomes of ecological research. NCEAS’s demonstrable successes (through two successful renewals, resulting in more than 15 years of continuous funding), combined with the continued quest for transformative research and the need to solve complex practical problems, have resulted in major national and international investments in synthesis. By 2017 nearly two dozen synthesis centers in various fields across the globe are based explicitly on the NCEAS model, representing public investments of many tens of millions of dollars (see e.g., http://synthesis-consortium.org/)

Synthesis centers vary in intellectual foundation and specific aims, but share similar purposes and operating principles, including: (1) a commitment to advance knowledge and address societal challenges through (2) small, self-organized collaborative groups of 6-20 scientists and practitioners (3) drawn from diverse disciplines, professional sectors, and social backgrounds (gender, nationality, seniority) whose work (4) combines spells of intensive, face-to-face collaboration in a setting insulated from day-to-day distraction and routine, separated by longer intervals of distal, computer-mediated work (Hackett et al. 2008; Palmer et al. 2016). Synthesis centers explicitly work to achieve the long-sought promise of interdisciplinary integration (Wilson, 1998; Sharp et al., 2011; NRC 2014).

Ethnographic studies of synthesis center collaborations identify several characteristics that might enhance interdisciplinary integration (Rhoten 2003; Hackett et al. 2008; Hackett and Parker 2016; Parker et al. 2018). Synthesis centers host concentrated collaborations in settings free from outside distractions and many of the usual marks of status (e.g., professor, student). Their small size and intense, immersive group dynamics mean that collaborators engage one another both as intellects and as whole persons. In turn, these qualities of group structure and interaction reduce status differences, balance participation, accelerate communication, and sustain trust, which allow ideas to be rapidly proposed, evaluated, and revised (Wooley et al. 2010; Hackett and Parker 2016; Bernstein et al. 2018). Synthesis centers are also resource-rich environments with full-time administrative and technical staff, resident researchers, and access to state-of-the-art computer software and hardware. Finally, synthesis centers instill a commitment to excellence among group members. They are ‘evocative environments’–places known to produce consequential research, challenging and motivating working group members to produce research of equal or greater quality (Zuckerman 1977). These are all beneficial aspects of synthesis centers and working group processes that are unlikely to be replicated in more traditional research environments, and which may help explain the remaining impact and influence associated with a paper originating in a synthesis center.

Synthesis working groups are formed by a scientific leader who develops a brief proposal to address a compelling scientific research question (often with direct implications for policy or practice) and identifies a group of 6-20 scientists and practitioners with distinctive and complementary expertise to work on the problem. Typically, groups may be formed, led, and composed of scientists from anywhere in the world. Proposals are competitively reviewed by a science advisory board. Working groups are diverse in composition, often including senior and junior scientists of various disciplines and specialties, as well as resource managers and environmental policy makers. The working group will gather at the center to work intensively for several days on several occasions over a period of 2 to 3 years, with group members remaining in communication with one another and working on aspects of their project during the intervals between meetings.

The immersive intensity of synthesis groups causes a distinctive pattern of social interaction that concentrates diverse expertise and promotes cooperation, collegiality, and transdisciplinary collaboration (that extends across academic disciplines to include government officials, decision makers, and representatives of civic groups. While these are primarily task-oriented groups, because they are immersive they also allow for shared leisure time, which may increase group cohesion and collegiality (Fine and Corte 2017). When conservation practice or policy is involved, as happens in about 25% of the groups, the consequences of the research become more visible and salient, lending focus, urgency, and excitement to the collaboration. For example, NCEAS research groups helped develop California’s Channel Islands Marine Protected Areas, informed the US Congress about honeybee decline, and studied the ecology of infectious diseases. In such cases the working groups included conservation or environmental policy experts, bringing into the collaboration the local concerns and practical needs of the particular site or problem (for example, species depletion in the Eastern Pacific fisheries or the ongoing stresses experienced by endangered species) and the distinctive perspective of creating knowledge that may provide a basis for intervention.

The structure and dynamics of synthesis collaborations may ease the challenges of interdisciplinary and cross-institutional research (Leahey et al. 2017; Cummings and Kiesler, 2008), and account for the exceptional quality and impact of research expected to emerge from synthesis centers. Several years of ethnographic observation (Hackett et al., 2008; Hackett and Parker, 2016), quantitative analyses of working group characteristics and performance (Hampton and Parker, 2010), and a pilot study using sociometric sensors (Parker et al., 2018) showed that synthesis center collaborations produced group characteristics that correspond to conditions that promote individual and collective creativity (Amabile, 2013; Parker and Corte 2017). These characteristics included: (1) resources, both in the form of human expertise and as research material and tools (including bridging social capital); (2) context, removed from everyday status cues and conducive to rich interpersonal interaction though bonding and shared social capital; (3) energy, arising from collective excitement about a motivating research question or compelling societal need (e.g., the use-inspired fundamental research of Pasteur’s Quadrant; Stokes 1997); and (4) adaptive management of ambivalence or values in tension. To illustrate, field observation revealed younger scientists speaking to senior scientists as equals, group bonding rituals and the development of distinctive identities and shared understandings, along with sharply critical interpersonal exchanges (which we called “peer review on the fly”) accelerating the creative process without rending the group, and rapid oscillation from constructive (brainstorming) modes of exchange to critical (evaluative review) of ideas, models, and data (Hackett and Parker 2016).

The same characteristics and dynamics observed in synthesis centers have manifested in other contexts that aim to inspire group synthesis. For example, Harvey (2014) studied Pixar, the animated film studio, and identified many of the same characteristics and dynamics observed in synthesis centers. Among those most conducive to creativity are resources (talent and technology), “a shared understanding that is unique to the collective” that holds the group together (Harvey, 2014: 325), and a process of construction and criticism much like peer review on the fly, in which “group members focus on single ideas in depth, ignore ideas, criticize ideas as they arise, and provide immediate interpersonal rewards for good ideas” (Harvey 2014: 328). In place of managing ambivalence or values in tension, Harvey conceptualizes creative synthesis as the product of a dialectical process.

Recent years have seen prominent and costly investments to build places, organize research, and shape group interactions to facilitate the integration of knowledge across disciplines (Kleinman et al. 2018). Examples include Stanford’s Clark Hall, which houses Bio-X, and interdisciplinary science and technology buildings on campuses as varied as Arizona State University, Northeastern University, and the University of Massachusetts, Amherst. The Howard Hughes Medical Institute built and operates the Janelia Research Campus to embody similar principles and goals. Concepts borrowed from synthesis centers, knowingly or not, inform the interdisciplinary collaborative initiatives of pioneering private foundations and patrons of science, including the Paul G. Allen Family Foundation, the Chan-Zuckerberg Initiative, and the cancer research investments of the Sean Parker Foundation.^5^ Synthesis centers have accelerated the development of collaborative communities, catalyzed research areas, and developed novel solutions to vexing societal concerns (Rodrigo et al. 2013; Palmer et al. 2016; Baron et al. 2017; Altschul et al. 2017).

Research examining the dynamics and performance of synthesis working groups has found that they spark distinctive and productive forms of social interaction, resulting in highly cited research and enduring career benefits for participants (Hampton and Parker 2011), yield effective solutions to socio-environmental problems (e.g., design of a successful marine protected area; Lubchenco et al., 2003), increase participants’ propensity to collaborate in the future (Rhoten and Parker, 2004), and enhance the likelihood of serendipitous and potentially transformative research (Hackett et al., 2008; Hackett and Parker, 2016).

Synthesis centers have altered the organization and conduct of research, but no analysis has yet addressed the fundamental question: *Do synthesis centers synthesize?* That is, first we ask if papers from synthesis centers integrate a greater diversity of topics than comparable papers from a reference corpus?^6^ We then ask if the topical diversity of a publication enhances its visibility or impact, as indicated by citations).

## What is diversity and why does it matter?

Diversity is a measure of the degree of difference within a collection of objects or ideas. We analyze diversity both as a composite concept (Rao-Stirling diversity) and in three aspects or dimensions--variety, evenness, and disparity-- each capturing a particular meaning of the concept (Rao 1982; Stirling 2007; Yegros et al., 2015).^7^ **Variety** is the number of different items present in a collection of objects or ideas (analogous to “species richness” in ecology): just as a more diverse or varied environment includes a greater number of species, a more diverse or varied publication would include a greater number of topics. **Evenness** is the relative frequency of occurrence of the items in a collection: a more diverse or even publication would include a more even (i.e., uniform, equal) distribution of topics. **Disparity** is the degree of difference between items: a more diverse or disparate publication includes topics that are less commonly associated with one another (or found together in a publication) and so are considered more *disparate* from one another. In short, a more diverse publication (and a more diverse literature) draws together a greater variety of topics, a more even distribution of these topics, with greater disparity between them (Patil and Taillie 1982; Stirling 2007; Yegros et al., 2015). We analyze diversity both as a composite measure and disaggregated into its dimensions.

Diversity matters because it indicates that ideas from different disciplines have been combined into a single publication. Science policymakers and program managers in foundations and federal funding agencies have encouraged interdisciplinary collaborations because their potential to recombine ideas in novel ways may yield innovative solutions to societal problems and original, potentially transformative, knowledge. The topical diversity of a publication is one indicator that interdisciplinary integration has occurred (NRC 2005; 2013).

## Hypotheses

If synthesis centers synthesize, then we expect their publications to be more diverse overall and more varied, balanced, and disparate than publications originating in other research environments. Synthesis center working groups are designed to include not only diverse disciplines but also stakeholders representing diverse sectors, such as government or the private sector, and the social organization and dynamics of synthesis center collaborations are designed to integrate the diverse ideas brought by participants to the collaboration. Synthesis centers bring together many fields of knowledge (variety) from disparate realms (disparity) in a balanced way (evenness), so we expect their publications to also manifest these dimensions of diversity. This leads to our first hypothesis:

> H1: *Synthesis papers display greater topical diversity than papers in the reference corpus.*

Size matters: larger collaborations may have greater breadth and depth, more network connections (social capital), greater credibility (cultural capital), and other advantages. Deliberately assembled to include the full range of expertise needed for a project, and generally funded well enough to include all necessary participants, synthesis collaborations are likely to be larger than the others. By virtue of such qualities, their greater size may make them also more diverse and make their article more visible. Thus, we also hypothesize that:

> H2: *Synthesis collaborations are larger than collaborations in the reference corpus.*

If synthesis collaborations truly differ from other collaborations in quality or character, as shown by the ethnographic studies described above, then such differences should express themselves as differences in diversity (aggregate and dimensions) that are not accounted for by differences in size (measured as numbers of authors, institutions, and references). Thus, we also hypothesize that:

> H3: *Size does not account for the greater diversity of synthesis center publications.*

Expectations are mixed about the influence of diversity and its dimensions on the visibility of publications and innovations (Fontana 2018). Research on innovation suggests that information pooled from disparate sources provides a foundation from which new ideas spring (Hargadon, 2002; Fleming and Waguespack, 2007). In the realm of science, some studies have found that articles and other scientific products (such as patents) that cover diverse topics have greater visibility (Shi et al., 2009; Schilling and Green, 2011; Uzzi et al., 2013; Leahey and Moody, 2014; Larivière, Haustein, and Börner, 2015; Lo and Kennedy, 2015; Leahey, Beckman, and Stanko 2017). Other studies suggest an inverted U relationship of visibility with increasing diversity (Larivière and Gingras, 2010; Yegros et al., 2015; Fontana et al., 2018). And Uzzi et al. (2013) found a more complex relationship with specific forms of diversity (a conventional knowledge base with only few atypical combinations) receiving the more visibility.

We contend that the heightened visibility (as gauged typically by citation counts) of synthesis center papers is not merely a function of the increased audience size that comes from covering more intellectual terrain (Leahey et al. 2017; Leahey & Moody 2014). Rather, papers that bring together and integrate ideas from disparate sources – that *synthesize* ideas – are more valued by the scientific community, and this explains their greater impact. These ideas drive our last two hypotheses:

> H4a: *Diverse papers (both in Rao-Stirling diversity and in each specific dimension) are more visible, even after controlling for collaboration size (authors, institutions, and references) and topic*
>
> H4b: *Synthesis center papers are more visible, even after controlling for diversity and its components (as well as collaboration size and journal impact factor)*

## Methods, Measures, and Analytic Approach

We test these hypotheses by using semantic analysis to compare the topical diversity of publications from synthesis centers (which we will call ‘synthesis papers’) with that of a reference corpus drawn from journals in cognate fields and from general science journals (which we call ‘reference papers’ or the ‘reference corpus’). By doing so, we focus the analysis on a measure of the substance or content of publications, rather than on characteristics of authorship groups (which we treat as an upstream property of a collaboration), social organization and dynamics (which we have studied in other work; Hackett and Parker 2016), productivity, or visibility (using citation-based measures, which we treat as a consequence of collaboration). Synthesis centers are represented by the two centers with the longest operational lives and publication records: NCEAS and the National Evolutionary Synthesis Center (NESCent) (1996-present and 2004-2014). We analyze words in the titles, abstracts, and keywords of publications to compare the topical diversity of peer-reviewed publications from NCEAS and NESCent with that of a reference corpus of publications representative of these fields (ecology and evolutionary biology, respectively).

We began with all articles published between 1997 and 2013 by scientists working at NCEAS (n=1213), and all articles published between 2004 and 2013 by scientists working at NESCent (n=335). These papers, totaling 1548 in all, were published in 112 different journals, and constitute our set of ‘synthesis papers.’ For comparison, we generated a reference corpus of literature that included 385,566 articles that appeared between 1997 and late 2013 in the 94 top journals (based on eigenfactor scores) for the five disciplinary areas most relevant to research done in NCEAS and NESCent (Ecology, Evolutionary Biology, Biodiversity Conservation, Fisheries, and Forestry). We also included articles from four general science journals *(Science, Nature, PLoS One,* and *PNAS),* and 14 journals that were common outlets for NCEAS and NESCent based research. Metadata for all articles were downloaded from the Web of Science.

To assess the diversity of ideas present in each article, we used Latent Dirichlet Allocation (LDA; Blei, Ng & Jordan 2003; Griffiths and Steyvers, 2003; DiMaggio et al. 2014) to construct and discover topics from the co-occurrence of words contained in article titles, abstracts, and keywords. LDA is an unsupervised probabilistic method of topic modeling that transforms the semantic content of documents into a proportional mixture of topics that is amenable to quantitative analysis. Topic modeling uses observed patterns of term co-occurrence within documents as a basis for probabilistic identification of latent ‘topics,” and then estimates the proportion of each document that is associated with each of the emergent topics. In contrast to classification schemes (such as Web of Science subject categories) or measures derived from an article’s bibliography, topic modeling offers a more detailed measure of the topic or substance of a published article, rather than focusing on its bibliographic ingredients (that is, characteristics of the references it cites). LDA’s ability to “generat[e] inductively classifications of ideas from texts” (Kaplan and Vakili 2015) offers a complementary method derived from substantive elements of publications.

LDA modeling requires setting initial parameters, such as the number of topics to be formed from the words in the corpus. Through experience, trial, and evaluation we settled on 154 substantive topics in the documents: this produced a set of topics that were neither too inclusive or general nor too specific.^8^

Using these topics, we calculated the Rao-Stirling Diversity index for each paper using this formula:

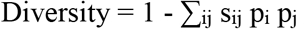

where p_i_ is the proportion of elements in topic “i,” p_j_ is the proportion of elements in topic “j,” and s_ij_ captures the degree of similarity between topics i and j, which we obtain from the LDA (Stirling 2007). To date, most applications of Rao-Stirling Diversity measures are based on topics derived from extant and fixed classification schemes, such as the Web of Science’s subject categories (Leahey, Beckman, and Stanko 2017; Yegros-Yegros et al. 2015), and such measures are usually applied to the bibliographic references of a paper—its ingredients—rather than to the semantic characteristics of the paper itself. Our use of LDA-derived topics as input for a diversity measure is novel. We also calculated and analyzed the three dimensions of diversity: variety (number of topics), evenness (the uniformity of the distribution of topics within an article, for a given number of topics), and disparity (the dissimilarity of the topics, given the number of topics) (Stirling 2007). These three dimensions are conceptually interrelated but only moderately correlated (see Appendix).

We found a small number of very distant outliers in the data, such as publications with more than 100 authors or references, which might bias the analysis. Therefore, for subsequent analysis, we truncated the distributions of addresses and references at the 99^th^ percentile to reduce their distorting influence; this is indicated by a “T” following the variable name. We also controled for topic (153 binary variables) and for other potential artifacts of the LDA approach.^9^ Finally, to ease comparisons in some analyses we standardized diversity variables to have mean = 0 and standard deviation = 1. This is indicated by a “Z” following a variable name.

To determine whether synthesis papers are not only more diverse but also (perhaps through their ability to synthesize such diversity) have higher visibility, we use a set of conventional measures and control variables. To measure visibility, we use the number of citations a paper has accrued as of 2013^10^ and a binary variable indicating whether or not the article is among the top 5% of cited articles. The binary variable focuses the analysis on the question of whether or not a contribution is a “hit” or a major contribution to its field (Uzzi et al. 2013; Lee et al. 2014).

Synthesis collaborations are designed to represent a breadth of scientific expertise and substantive knowledge, and have funds to assemble such groups, and so they may have more members than others. Greater size, in turn, brings not only expertise and knowledge but also various other forms of human, social, and cultural capital (Collins 2000, Simonton 2004; Burt 2005; Lee et al. 2015; Wuchty et al. 2007; Uzzi et al. 2013; Leahey et al. 2017). Meta-analyses conducted by synthesis center groups have twice as many authors, studied 1.6 times as many species, referenced 1.4 times as many publications, analyzed 1.3 times as many datasets, and were published in higher impact journals compared to meta-analyses that originated in places other than synthesis centers (Cadotte et al., 2012).

We take size into account with three variables: the number of authors of a paper, an indicator of the size of the collaboration; the number of distinct institutions (addresse) represented by authors, which is an indicator of substantive breadth and social capital (Burt 2005); and the number of references cited in an article, which is an indicator of the breadth of an article’s intellectual foundation (a form of cultural capital; Collins 2000; Simonton 2004). Each of these – individuals, organizations, and references – is an intellectual resource that may contribute to the diversity and visibility of an article.^11^ We hypothesize that these characteristics of the collaboration not only may account for differences in diversity and visibility, but also may play a mediating role through which properties of synthesis center collaborations influence article diversity and visibility.

## Results

The Rao-Stirling diversity value for synthesis center papers is virtually identical to that of papers in the reference corpus. But comparing mean values for each of the three dimensions of diversity yields a more complicated result: synthesis center papers include more topics than papers in the reference corpus (that is, have greater variety), and the distribution of terms among topics in synthesis papers is more even than the distribution in reference corpus papers (that is, have greater balance), lending support to Hypothesis 1. Somewhat surprisingly, synthesis center papers have significantly *lower* disparity (measured as cosine similarity) than the reference corpus, suggesting that they include topics that are more closely related to one another. We examine this finding in greater detail below.

Collaboration size does not appear to mediate the relationship between synthesis center affiliation and diversity. On average, synthesis center papers have slightly (but not significantly) larger authorship groups than papers in the reference corpus, and significantly greater numbers of institutional affiliations and references (see Table 2), lending partial support to Hypothesis 2. Even when year of publication and modal topic are controlled (Table 3), synthesis center papers have significantly more authors from more institutions and cite more literature than papers in the reference corpus. However, contrary to Hypothesis 3, we find that papers with more authors are not more diverse (see Table 4): in fact, on all measures, a larger authorship group is associated with less diversity (both overall and in the variety and disparity components). Thus, synthesis center collaborations are more diverse, but this effect cannot be explained by their larger size (i.e., numbers of authors, institutions, and references). Even after controlling for other variables^12^, the differences in Table 1 remain: publications of synthesis center collaborations have greater variety and balance (but not disparity), and now also have greater Rao-Stirling diversity, than do papers in the reference corpus (Table 4). The effect of synthesis center affiliation is substantial: its effect on variety is equivalent to adding five authors to a paper, and its effects on evenness is equivalent to adding a dozen institutions.

**Table 1.** Mean Differences in Diversity and Its Components, Synthesis Centers and Reference Corpus

|  | A | B | C | D | E |
| --- | --- | --- | --- | --- | --- |
| Variable | Synthesis center articles | Reference corpus | Difference (t) | Mean (SD) | N |
| Diversity (Rao-Stirling) | .277 | .277 | .000 (.004) | .277 (.123) | 385,565 |
| Variety | 6.75 | 6.18 | .578 (10.3)*** | 6.18 (2.17) | 383,834 |
| Balance | .855 | .812 | .043 (13.3)*** | .812 (.127) | 383,835 |
| Disparity | .417 | .461 | -.043 (11.7)*** | .461 (.145) | 383,835 |
| N | 1883 | 383,653 |  |  |  |

**Table 2.** Mean Differences in Collaboration and Publication Characteristics, Synthesis Centers and Reference Corpus

| Variable | A<br>Synthesis<br>center<br>articles | B<br>Reference<br>corpus | C<br>Difference (t) |
| --- | --- | --- | --- |
| Number of authors | 5.19 | 4.97 | .23 (.980) |
| Number of institutions | 4.48 | 2.83 | 1.66 (22.2)*** |
| Number of references | 56.0 | 44.4 | 11.6 (21.8)*** |
| Proportion of topic<br>weight removed as junk | .354 | .334 | .021 (8.31)*** |
| Count of tokens per<br>document (after stop<br>word removal) | 109.3 | 111.3 | 2.00 (2.35) |
| Publication year | 2006.0 | 2006.8 | .737 (6.03)*** |

N=385,566
\* = $p < .05$ ; \*\* = $p < .01$ ; \*\*\* = $p < .001$

**Table 3.** OLS Regression of Numbers of Authors, Institutions, and References on Synthesis Center Affiliation and Control Variables

|  | A | B | C | D | E |
| --- | --- | --- | --- | --- | --- |
|  | Number of<br>Authors | Number of<br>Institutions | Number of<br>References<sup>10</sup> | Institutions<sup>T</sup> | References<sup>T</sup> |
| Synthesis<br>center paper<br>(1=yes) | .890*** | 1.42*** | .777*** | 1.01*** | .773*** |
| Number of<br>Authors | -- | -- | -- | .470*** | .004** |
| Constant | 2.86 | 4.01 | 5.62 | 2.67 | 5.60 |
| R<sup>2</sup> adj | .27 | .13 | .15 | .37 | .15 |
| N | 385,566 | 385,566 | 385,566 | 385,566 | 385,566 |

Note: Although coefficients are not shown, we also controlled for year of publication and modal topic (indicator variables for 153 topics). \* = $p < .05$ ; \*\* = $p < .01$ ; \*\*\* = $p < .001$

**Table 4.** OLS Regression of Diversity and Components on Synthesis Center Origin, Collaboration Characteristics, and Control Variables

|  | A | B | C | D |
| --- | --- | --- | --- | --- |
|  | DiversityZ | VarietyZ | BalanceZ | DisparityZ |
| AuthorsT | -.018*** | -.010*** | -.001 | -.004*** |
| InstitutionsT | .017*** | .010*** | .002** | .004*** |
| References10 | -.013*** | .112*** | .003*** | -.028*** |
| VarietyZ |  | -- | .537*** | .460*** |
| BalanceZ |  | .474*** | -- | -.238*** |
| DisparityZ |  | .353*** | -.206*** | -- |
| Synthesis center paper (1 = yes) | .077** | .056*** | .050** | -.078*** |
| Constant | .104 | .796 | -1.02 | .167 |
| R2 | .11 | .44 | .38 | .28 |
| n | 385,566 | 383,835 | 383,835 | 383,835 |

Note: All models also control for year of publication, topic weight removed, number of tokens, and modal topic (dummy variable with 153 categories).
\* = $p < .05$ ; \*\* = $p < .01$ ; \*\*\* = $p < .001$

We expected that papers from synthesis centers would be more diverse in every respect, but instead find that they have less disparity (that is, greater similarity) than papers in the reference corpus (see Table 4, column D). This is particularly surprising considering the various measures of size and variety (authors, addresses, references) that favor synthesis center papers. Below we discuss this result and its possible roots in the distinctive structure and interaction patterns of synthesis center collaborations.

As expected (Hypotheses 4b), synthesis center papers garner more citations than papers in the reference corpus. As shown in Table 5, the differences are substantial: twice as many citations and twice the probability of being among the top 5% (“hits” or very visible articles). But recall that synthesis collaborations are larger in some respects (institutions, references; see Table 4) and the publications they produce are more diverse than those of the reference corpus, so it is necessary to consider size and diversity, and to include other variables, to determine whether (and the means by which) synthesis collaborations produce more visible publications.

**Table 5.** Means of Publication Impact Measures for Synthesis Papers and Reference Corpus

|  | A | B | C |
| --- | --- | --- | --- |
| Variable | Synthesis center articles | Reference corpus | Difference (t) |
| Citations received (as of 2013) | 82.7 | 41.0 | 41.6 (11.9)*** |
| Fraction in top 5% of all articles | 11.9% | 5.3% | X2 128.9*** |

Significance levels indicated by asterisks: \* = $p < .05$ ; \*\* = $p < .01$ ; \*\*\* = $p < .001$

These differences in visibility are not explained by group size or diversity (in the aggregate and by dimensions). To determine this, we modeled two visibility outcome variables: 1) the number of citations, using negative binomial regression for count data, and 2) the binary property of a paper being a “hit” (in the top 5% of the citation distribution) or not, using a logistic regression model. Control variables in each model include the three dimensions of diversity (variety, balance, and disparity), size (number of authors, institutions, and references), and synthesis center affiliation. Table 6 (columns A and C) show that synthesis center collaboration has a strong positive effect on both outcomes, even after controlling for size and diversity.

**Table 6.** Regression of Measures of Publication Visibility on Synthesis Center Affiliation and diversity measures, Variety, Balance, and Disparity, Collaboration Characteristics, and Control Variables

|  | A | B | C | D |
| --- | --- | --- | --- | --- |
|  | Article citations | Article citations | Top 5% (1=yes) | Top 5% (1 = yes) |
| AuthorsT | .062*** | .062*** | .189*** | .189*** |
| InstitutionsT | .017*** | .044*** | .056*** | .056*** |
| References10 | .116*** | .096*** | .130*** | .130*** |
| VarietyZ | -.040*** | -.060*** | -.074*** | -.076 |
| BalanceZ | -.013*** | -.043*** | -.172*** | -.172 |
| DisparityZ | -.025*** | -.039*** | -.070*** | -.070 |
| Synthesis center paper (1 = yes) | .506*** | .586*** | 1.20*** | 1.03*** |
| VarietyZ*synth |  | .059** |  | .154* |
| BalanceZ*synth |  | .067** |  | .245* |
| DisparityZ*synth |  | .007 |  | -.085 |
| Influence |  |  |  |  |
| Constant (alpha) | .758 | 3.85 | -2.48 | -2.48 |
| n | 383,835 | 383,835 | 353,783 | 353,783 |
| Pseudo R2 | .15 | .12 | .34 | .34 |

Note: all models also control for: year of publication; topic weight removed, number of tokens, and modal topic (dummy variable with 153 categories).
For all predictor variables except moderator effect \* = $p < .05$ ; \*\* = $p < .01$ ; \*\*\* = $p < .001$ ; for moderator effects p levels are reported for one-tailed tests because specific directional hypotheses were proposed: \* = $p \leq .10$ . \*\* = $p \leq .05$ , \*\*\* = $p \leq .01$ .

Given previous literature, it is surprising that diversity and its components have significant *negative* effects on both measures of visibility. Indeed, H4a expects a positive effect. We acknowledge that the difference may be a consequence of our use of topic modeling (versus bibliometric measures of interdisciplinary research), particular control variables used in the models, or other such differences in method. That said, this result suggests that unmeasured characteristics of synthesis collaborations affect the topical diversity of publications in ways that increase their visibility. To explore this possibility, we examine whether synthesis center affiliation moderates the influence of diversity on visibility. Moderation would mean that the effects of diversity or one of its components is different for synthesis papers than it is for papers in the reference corpus. Models B and D in Table 6 include interaction terms that evaluate this possibility. The results suggest that there is a moderating effect: for synthesis center papers only, variety and balance (but not disparity) have positive and marginally significant effects on citation count and on the binary “top 5%” or “hit” variable (p < .10, one-tailed test). Collaborations organized and conducted under the auspices of synthesis centers appear to have qualities that convert the potential liabilities of diversity and its components into assets, overcoming the liability of recombinatory innovation (Hampton and Parker 2011; Kaplan and Vakili 2015; Hackett and Parker 2016). This contingent result is similar to Uzzi et al (2013), who found that just enough disparity, not too much, helps a paper, and with Lee et al. (2014), who found that variety interacted with size to influence the visibility of an article.

## Summary and Discussion

Our comparison of synthesis center papers and a reference corpus reveals that synthesis center papers are more diverse (in terms of the number of topics integrated, and the evenness of those topics), and that these differences remain when control variables are included in the models. Differences in the size and the social and cultural capital of authorship groups partly mediate the effect of synthesis center affiliation on diversity. For the reference corpus, the components of diversity (variety, balance, and disparity) are associated with *lower* influence and citation scores, but for synthesis center papers diversity brings *greater* influence and citations. This holds true as a simple difference in means and in multivariate models that control for dimensions of diversity, characteristics of authorship teams, and other variables.

Do synthesis centers synthesize by bringing together diverse topics into a single publication? If diversity is measured solely by mean difference in the Rao-Stirling (aggregate) measure of diversity, then the answer is no (Table 1). But diversity is a complex concept, and disaggregating it into three components (variety, evenness, and disparity) reveals that synthesis centers produce papers with significantly greater variety and balance, but significantly lower disparity, than papers in the reference corpus. This is mediated by the greater size of synthesis center collaborations (numbers of authors, distinct institutions, and references, which may stand for their greater social and cultural capital; Table 2). To some degree, size may play a mediating role, but even when taking size into account, there is a persistent direct effect: synthesis center papers have significantly greater variety and balance, but less disparity, than papers in the reference corpus (Table 4).

It is somewhat surprising to find that large authorship groups are associated with lower levels of diversity. If the main purpose of collaboration is to pool specialized knowledge (Maienschein 1993; Hackett 2005), then papers with more authors should have greater diversity. But perhaps that is not the dominant motivation for collaboration (Leahey and Reikowsky 2008; Leahey 2016). Perhaps collaboration occurs to add person-power to accomplish a shared set of similar tasks, rather than a differentiated set of dissimilar tasks. Or, perhaps, a topic (in the sense of this paper) is broader than a scientist’s expertise and so two or more scientists may be needed to accomplish the work contained within a topic. And, finally, the causal arrow may run in the opposite direction: perhaps a substantial degree of intellectual and interpersonal *similarity* is necessary to sustain and hold together a collaboration with many members (cf. Farrell 2001; Parker and Corte 2017). Components of the complex concept “size” and components of the complex concept “diversity” may have distinctive relationships with one another. For example, Table 4 shows that the number of institutions in a collaboration significantly increases overall diversity and all its components, but that the number of references (an indicator of size that emphasizes intellectual or cultural capital) increases two components of diversity--variety and balance—but not the third (disparity).

Others who have examined the effects of organizational diversity on scientific performance have also found inconsistent results. Cummings and Kiesler’s (2005) analysis of 491 multi-university collaborations found that the number of institutions involved in a collaboration was the single strongest predictor of *lower* collaborative success. In contrast, Parker and Hampton (2011) found that working groups with a higher ratio of institutions to members performed better. The difference may be that synthesis center collaborations include extended periods of intense face-to-face interactions, which improves coordination among collaborators, enhances communication, and builds trust, solidarity, and commitment that sustain the group through periods of remote collaboration (Collins 1998; Hackett and Parker 2016). Recurrent face-to-face meetings also affords groups time to surface and resolve differences, producing an agreed-upon central message.

Synthesis center papers are more visible than papers in the reference corpus (Table 5), and such difference are mediated, in part, by size and diversity dimensions (Table 6). While such qualities of collaboration partly account for differences between synthesis papers and the reference corpus, with such variables controlled synthesis center papers still have significantly (and substantially) greater influence and citation counts than papers in the reference corpus (Table 6). Synthesis center effects are mediated, to some degree, by collaboration size and the dimensions of diversity, and the effects of those components are moderated, to a modest degree, by synthesis centers. But the strong and significant positive effect of synthesis centers on article influence and citation count remain to be explained.

## Conclusion

Scientific synthesis has arisen rapidly in response to challenges such as overcoming hyper-specialization, navigating immense and growing literatures, conceptualizing complex socio-environmental problems, and enhancing the potential for serendipitous discovery and transformative research. Synthesis is essential in a world where scientific specialists must collaborate to solve complex intellectual puzzles and ‘wicked’ practical problems that lie beyond the reach of any one discipline, profession, dataset, method, or theory. Research has shown the synthesis working group to be a distinctive form of scientific collaboration that reliably produces consequential, high-impact publications, but no one has attempted to directly investigate their *raison d’être:* do synthesis working groups produce publications that are substantially more diverse than those produced outside of synthesis centers, and if so, how and with what effects? We have investigated these questions through a novel textual analysis. Let us emphasize that this is a novel approach: We are not sure how measuring diversity in terms of topics obtained from topic modelling rather than from co-citation, bibliographic coupling, Web of Science categories, or other bibliometric means, though we do know that such measures often disagree with one another (Wang and Schneider 2018). We have not tested the measure by validating it against other properties of specific articles or researchers, but we do know that it taps into the substance of the articles—words—rather than more distal properties. The power and robustness of the measure remain to be determined, yet we think it has shown sufficient promise to merit further investigation as a complement to bibliometric approaches. What have we learned?

Overall, synthesis center publications have greater numbers of authors from more institutions than do publications in the reference corpus, and these integrate a broader conceptual and knowledge base, as measured by numbers of references. Surprisingly, having greater numbers of authors is not associated with greater topical diversity, but having a greater number of distinct institutional addresses did increase diversity. Papers with more authors and more institutions are also more highly cited; article diversity, as a whole or in components, is strongly and negatively related to citation counts and the probability of being a hit paper (i.e., falling within the top 5% of the citation distribution). Synthesis papers are more topically diverse, highly cited, and influential, suggesting that unmeasured properties of the synthesis center collaboration are responsible for the differences.

Our research also yields several practical lessons. First, the positive association of synthesis center papers with diversity, citations, and influence strongly suggests that despite the current excitement around ‘virtual organizations’ and distal forms of collaboration, there is still a place for physical centers and face-to-face groups. They remain our best opportunity to produce transformative and synthetic research. Second, policies intended to identify and support transformative research (NSB 2008) have attempted to do so by selecting particularly promising projects or people, generally with very low award rates. This study suggests that there is merit in creating organizations, such as synthesis centers, that integrate diverse concepts, methods, and data. Third, text analysis is a rapidly evolving field with substantial promise for revealing the substance and intellectual dynamics of science, complementing bibliometric measures of scientific properties and performance. Finally, current demands for transformative scientific knowledge and innovative solutions to pressing practical problems have stimulated policy and programmatic interest in convergence (Sharp et al. 2011; NAS 2014). Such organizational innovations are in their infancy and should be regarded as experiments, informed and adaptively managed by analyses of their collaborative processes and research outcomes.

## Acknowledgements

We thank Stacy Rebich Hespanha for her expert assistance with the LDA and data set preparation. This work was supported by NSF grant SBE1242749 to Ed Hackett and John Parker, and group meetings were graciously hosted and travel paid by NCEAS and NESCent.

## Appendix

Correlations among Diversity and Components
|  | Variety | Balance | Disparity | Diversity |
| --- | --- | --- | --- | --- |
| Variety |  |  |  |  |
| Balance | .469 |  |  |  |
| Disparity | .277 | -.094 |  |  |
| Diversity | .644 | .479 | .676 |  |

Correlations among size measures
|  | Authors | Addresses | References |
| --- | --- | --- | --- |
| Authors |  |  |  |
| Addresses | .645 |  |  |
| References | -.004 | .036 |  |

N = 387,040
P < .02 for all correlations

## Footnotes

1 Following other analyses of interdisciplinarity, ours is “based on the concept of integration: a mode of research that integrates concepts or theories, tools or techniques, information or data from different bodies of knowledge” (NRC 2005; Yegros et al. 2015: 7). This definition is more than convenient: it also invokes the conceptualization of creativity as grounded in the association of different ideas (Mednick 1962; Amabile 1983; Simonton 2004).

2 Consilience, a process of “jumping together” (jumping is the “*siliens*” part, as in resilience), proposes that diverse fields of knowledge—not just sciences but also humanities and social sciences—would jump together through an almost elective affinity to address complex societal and intellectual problems with broader, deeper, and more fundamental (some might say mechanistic, even bio-reductionist) explanations. Convergence asserts that certain fields are bending, turning, tending toward one another (the literal meaning of the Latin root *verger*), and perhaps need some assistance (or removal of resistance) to accelerate the process. For example, an MIT report (Sharp 2011) argued for investing in the convergence of the life sciences and engineering to bring fundamental knowledge coupled with know-how to bear on health needs (and, reciprocally, to use health needs to inspire fundamental research). NAS (2014) issued a report on convergence, and convergence is among the 10 Big Ideas guiding NSF investments (https://www.nsf.gov/news/special_reports/big_ideas/).

3 Other US centers focused on evolutionary theory and mathematical biology. About 24 synthesis centers have been developed worldwide, across all fields of science.

4 Other analyses use the co-occurrence of subject matter categories of the references in a paper to measure its diversity (Porter et al. 2007; Uzzi et al., 2013; Yegros et al. 2015). We think the words in the title, abstract, and keywords offer a complementary view of interdisciplinarity that is based on the output of research rather than the ingredients.

5 (https://www.alleninstitute.org/what-we-do/frontiers-group/; http://parker.org/about; https://chanzuckerberg.com/)

6 Unlike measures of interdisciplinarity that are applied to the bibliographic references of an article—its ingredients--topical diversity is an indicator applied to an intellectual product (in this case a published article).

7 Rao-Stirling is one of a family of diversity measures, known as Leinster-Cobbold diversity (Mugabushaka, Kyriakou, and Papazoglou, (2016).

8 LDA is substantively naïve and so, along with meaningful topics the method also creates a small number of topics that convey little substantive meaning about the paper, formed by the co-occurrence of numeral (one, two, three), directional (up, down), and comparatives (greater, lesser) terms. When such terms co-occur they create an apparent topic with no clear substantive meaning that we treated as a data artifact, as is usual practice (Kaplan and Vakili 2014). We removed such terms from the analysis and used the fraction of a topic’s weight that was removed in this fashion as a control variable in our analysis to account for any effect this may have had on outcomes of interest here. For similar reasons we also controlled for differences in the number of valid words or multi-word terms for each paper. Weighted sets of representative terms for each topic give substantive meaning to topics, and a solution that yielded 154 substantive topics (and 46 meaningless topics) was judged most representative of the substance of the papers.

9 These other control variables are percent of topics removed and number of tokens used to characterize an article.

10 We recognize that citations are not always positive (see MacRoberts and MacRoberts 1996). However, citations to work indicate its usefulness and provide visibility in the scientific community – both of which signal impact.

11 Numbers of authors and institutions are correlated .645; number of references is almost uncorrelated with authors and institutions; see Appendix.

12 Other qualities of collaborations that we have not measured here (but have studied with other methods and reported elsewhere; see Hackett and Parker 2016; Hampton and Parker 2011) may also influence diversity. For example, how much group members have worked together outside this particular collaboration, or group leaders’ selection biases.

